# The fungal expel of 5-fluorocytosine derived fluoropyrimidines mitigates its antifungal activity and generates a cytotoxic environment

**DOI:** 10.1101/2022.08.24.504767

**Authors:** Luis Enrique Sastré-Velásquez, Alex Dallemulle, Alexander Kühbacher, Clara Baldin, Laura Alcazar-Fuoli, Anna Niedrig, Christoph Müller, Fabio Gsaller

## Abstract

Invasive aspergillosis remains one of the most devastating fungal diseases and is predominantly linked to infections caused by the opportunistic human mold pathogen *Aspergillus fumigatus*. Major treatment regimens for the disease comprise the administration of antifungals belonging to the azole, polyene and echinocandin drug class. The prodrug 5-fluorocytosine (5FC), which is the only representative of a fourth class, the nucleobase analogs, shows unsatisfactory *in vitro* activities and is barely used for the treatment of aspergillosis. The main route of 5FC activation in *A. fumigatus* comprises its deamination into 5-fluorouracil (5FU) by FcyA, which is followed by Uprt-mediated 5FU phosphoribosylation into 5-fluorouridine monophosphate (5FUMP). In this study, we characterized and examined the role of a metabolic bypass that generates this nucleotide *via* 5-fluorouridine (5FUR) through uridine phosphorylase and uridine kinase activities. Resistance profiling of mutants lacking distinct pyrimidine salvage activities suggested a minor contribution of the alternative route in 5FUMP formation. We further analyzed the contribution of drug efflux in 5FC tolerance and found that *A. fumigatus* cells exposed to 5FC reduce intracellular fluoropyrimidine levels through their export into the environment. This release, which was particularly high in mutants lacking Uprt, generates a toxic environment for cytosine deaminase lacking mutants as well as mammalian cells. Employing the broad-spectrum fungal efflux pump inhibitor clorgyline, we demonstrate synergistic properties of this compound in combination with 5FC, 5FU as well as 5FUR.

## Introduction

Although several fungal species are able to cause life-threatening disease types, the highest death tolls are linked to infections by members of the genera *Aspergillus, Cryptococcus, Candida* and *Pneumocystis* [1, 2]. Among pathogenic *Aspergilli,* the major cause of life-threatening invasive aspergillosis remains the most prevalent mold pathogen *Aspergillus fumigatus* [3]. Being constantly exposed to its airborne spores, humans can inhale several hundred spores daily, which, due to their small size, are capable of entering the lung alveoli [4]. While the immune system in healthy individuals readily eliminates spores, their entrance in the lung is particularly dangerous for patients with a defective immune defense [4–6]. Widely employed treatment regimens against aspergillosis involve the use of azole, polyene or echinocandin antifungal agents [7, 8]. The nucleobase analog 5-fluorocytosine (5FC) is barely used to treat *Aspergillus*-induced diseases [7]. One obvious reason represents its negligible activity *in vitro,* however, previous work demonstrated a beneficial outcome of 5FC treatment on survival in murine infection models [9, 10]. In contrast to *Aspergillus,* 5FC shows potent activity against a variety of pathogenic species belonging to the genera *Candida* and *Cryptococcus.* For the treatment of infections caused by these pathogens, 5FC monotherapy is typically avoided due to rapid occurrence of resistance [11].

Most likely due to their apparent requirement for 5FC activity, commonly detected mechanisms of resistance in yeast were connected to mutations in genes that impair its uptake *(FCY2; A. fumigatus fcyB*), cytosine deaminase *(FCY1; A. fumigatus fcyA)* and uracil phosphoribosyltransferase (*FUR1*; *A. fumigatus uprt*) [11]. In addition to these, mutations leading to increased pyrimidine production as well as other genetic factors appear to play important roles in 5FC resistance [11]. In *Cryptococcus gatti*, for instance, mutations in the UDP-glucuronate decarboxylase (*UXS1*) gene increased 5FC resistance. Defects in the enzyme elevated glucuronic acid levels and led to changes in nucleotide metabolism [12]. In *Candida glabrata* an important role of aquaglyceroporins *(FPS1* and *FPS2)* in 5FC tolerance was suggested. Inactivating their gene function elevated 5FC intracellular accumulation, which resulted in increased 5FC susceptibility [13]. In *A. fumigatus,* a clear dependency of 5FC activity on the environmental pH has been demonstrated [9, 14]. At neutral pH, which is the common pH for antifungal susceptibility testing [15, 16], this mold displays intrinsic 5FC resistance, which was linked to transcriptional repression of *fcyB*, coding for the main 5FC uptake protein [14]. At acidic pH, 5FC activity is significantly enhanced by upregulation of the gene [14].

In contrast to azoles, polyenes and echinocandins, 5FC per se is not toxic. For its activation, 5FC requires conversion into the nucleotide 5-fluorouridine monophosphate (5FUMP). Subsequent to its synthesis, 5FUMP is further metabolized into active 5-fluorinated ribo- and deoxyribo-nucleotides that hamper cell growth [11]. The main route of 5FUMP formation in fungi involves two enzymatic steps including 5FC deamination into 5FU by cytosine deaminase and phosphoribosylation thereof into 5FUMP *via* uracil phosphoribosyltransferase (**Fig 1A**) [11]. Compared to fungal species that are sensitive to 5FC, mammalian cells lack cytosine deaminase but possess two main routes of 5FUMP formation out of 5FU. 5FU can either be directly converted into 5FUMP by the *de novo* pyrimidine biosynthesis enzyme orotate phosphoribosyltransferase or indirectly *via* 5-fluorouridine (5FUR), which requires the consecutive action of uridine phosphorylase and uridine kinase [17].

**Fig 1.**
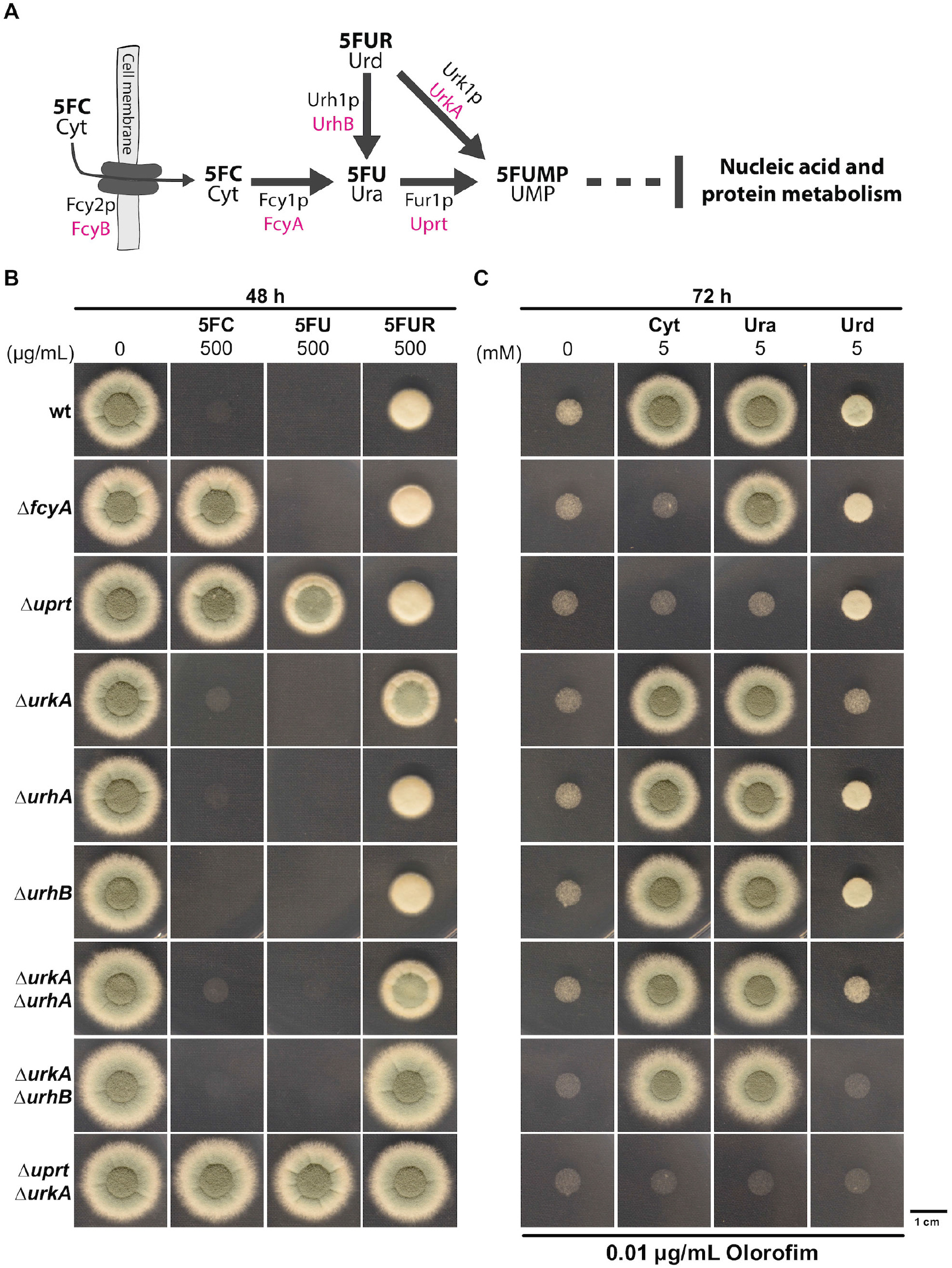
UrhB and UrkA mediate the hydrolysis and phosphorylation of 5FUR/uridine. **(A)** Scheme illustrating proteins involved in 5FC uptake and pyrimidine salvage pathway based metabolization of fluoropyrimidines in yeast (black) and *A. fumigatus* (magenta). **(B)** The role of uridine nucleosidase and uridine kinase encoding genes was determined by susceptibility testing. **(C)** Resistance analysis was complemented with a reversal assay studying the contribution of enzymes in the metabolization of nucleobases cytosine (Cyt) and uracil (Ura) as well as the nucleoside uridine (Urd) during *de novo* pyrimidine biosynthesis inhibition.

Here, we characterized further pyrimidine salvage enzymes in *A. fumigatus* that participate in 5FC metabolic activation and demonstrate that, like mammalian cells, this mold harbors the enzymatic repertoire to generate 5FUMP indirectly. Measurements of extracellular 5FU and 5FUR during 5FC exposure indicated that *A. fumigatus* cells export both 5FC derivatives into the environment, which alleviates 5FC activity and, simultaneously, creates a cytotoxic environment. Drug interaction analysis of 5FC, 5FU and 5FUR employing a fungal efflux pump inhibitor [18, 19] revealed synergistic properties with each pyrimidine analog and suggests that 5FC antifungal activity could be enhanced when used in combination with fungal efflux pump inhibitors.

## Results

### *Aspergillus fumigatus* possesses pyrimidine salvage activities that mediate hydrolysis and phosphorylation of 5FUR

The pyrimidine salvage enzymes cytosine deaminase (FcyA, AFUB_005410) and uracil phosphoribosyltransferase (Uprt, AFUB_053020) are key enzymes in 5FC metabolic activation in *A. fumigatus.* While loss of FcyA (Δ*fcyA)* only confers hyper-resistance to 5FC, the absence of Uprt (Δ*uprt*) leads to high levels of resistance to 5FC and 5FU (≥ 100 μg/mL) [20]. In this work, resistance profiles of Δ*fcyA* and Δ*uprt* have been further assessed using 5FC and 5FU hyper doses and found that in the presence of 500 μg/mL 5FU, growth of Δ*uprt* was clearly inhibited (**Fig 1B**). This indicated the presence of further enzymes that are involved in 5FC metabolism.

Previous work in yeast demonstrated the presence of genes encoding pyrimidine salvage enzymes with uridine nucleosidase (Urh1p) and uridine kinase (Urk1p) activity mediating/with their main functions being the hydrolysis of uridine to uracil and the phosphorylation of uridine into uridine monophosphate, respectively [21]. BLASTP analysis using the respective *S. cerevisiae* protein sequences suggested the presence of a putative Urk1p ortholog, here named UrkA (AFUB_022460) as well as two Urh1p orthologs in *A. fumigatus,* here termed UrhA (AFUB_005870) and UrhB (AFUB_011230) (**S1A** and **S1B Table**). To analyze the role of these genes in 5FUR/uridine metabolization, coding sequences of *urkA, urhA* and *urhB* were deleted in wild-type (wt). In addition, double deletion mutants Δ*urkA*Δ*urhA* and Δ*urkA*Δ*urhB* were generated, anticipating combined loss of 5FUR/uridine hydrolysis and phosphorylation activities. To further test that 5FUMP formation can only occur through Uprt or UrkA, the double deletion mutant Δ*uprt*Δ*urkA* was made. This mutant still encodes activities enabling the hydrolysis of 5FUR/uridine into 5FU/uracil and *vice versa,* however, the formation of 5FUMP/UMP should be blocked. The individual gene functions were analyzed following two approaches. One included 5FC, 5FU and 5FUR resistance analysis of the different salvage pathway mutants. The second aimed to investigate the capacity of those strains to utilize cytosine, uracil and uridine for UMP generation during simultaneous inhibition of its *de novo* biosynthesis. In this case, the antifungal drug olorofim, inhibiting the essential enzyme dihydroorotate dehydrogenase [22], was supplemented to the medium.

Single mutants Δ*urhA* and Δ*urhB* showed wt-like susceptibility to 5FC, 5FU and 5FUR and growth of these strains could be promoted during olorofim treatment with either cytosine, uracil or uridine (**Fig 1B** and **1C**). In agreement with the role of UrkA in the formation of 5FUMP/UMP out of 5FUR/uridine, Δ*urkA* displayed increased resistance to 5FUR and its growth could only marginally be recovered by uridine supplementation during *de novo* pyrimidine biosynthesis inhibition. This further indicates that 5FUR/uridine hydrolysis coupled to Uprt mediated 5FU/uracil phosphoribosylation is less efficient than the direct conversion of 5FUR/uridine into 5FUMP/UMP. Surprisingly, while Δ*urkAΔurhA* with predicted block in 5FUR/uridine phosphorylation and hydrolyzation phenocopied the single deletion mutant Δ*urkA* with regard to its 5FUR resistance, Δ*urkA*Δ*urhB* displayed 5FUR hyper-resistance. By reintroducing a functional gene copy of *urhB* into Δ*urkA*Δ*urhB*, 5FUR resistance levels were restored, similar to that of Δ*urkA* (**S1 Fig**). This suggests that despite the higher homology of UrhA to yeast Urh1p (**S1A Table**), in *A. fumigatus* the hydrolysis of 5FUR/uridine is mainly mediated by UrhB. Δ*uprt*Δ*urkA* lacking both enzymes for direct 5FUMP/UMP generation, displayed hyper-resistance to 5FC, 5FU and 5FUR (**Fig 1B**) and its growth could not be promoted by the addition of cytosine, uracil or uridine to the growth medium (**Fig 1C**). Interestingly, despite the potential presence of salvage activities that enable the formation of 5FUMP/UMP independently of Uprt, growth of Δ*uprt* could not be recovered by cytosine or uracil during olorofim treatment. This, together with the high degree of 5FU resistance observed for Δ*uprt* compared to Δ*urkA,* suggests that 5FUMP/UMP synthesis occurs predominantly *via* Uprt.

### *A. fumigatus* has the enzymatic capacity to convert 5FU into 5FUR

The growth impairment of Δ*uprt* at high concentrations of 5FU that can be overcome by the additional inactivation of *urkA* (Δ*uprt*Δ*urkA*) (**Fig 1B**), strongly suggests that *A. fumigatus* is able to convert 5FU into 5FUR. The catalytic cleavage of uridine into uracil by uridine nucleosidase (EC: 3.2.2.3) Urh1p, however, is anticipated to be irreversible [23]. Therefore, the enzyme that generates 5FUR out of 5FU in *A. fumigatus* still remained unclear. To rule out that *A. fumigatus* UrhA and/or UrhB might be involved in the conversion of 5FU into 5FUR, we first generated Δ*uprt*Δ*urhA* and Δ*uprt*Δ*urhB* double mutants and compared 5FU resistance to that of the recipient strain Δ*uprt.* Both double mutants showed the same level of 5FU resistance as their progenitor strain Δ*uprt* (**Fig 2A**), which suggested that the hydrolysis of 5FUR by uridine nucleosidase also occurs irreversible in *A. fumigatus.* In bacteria and mammals an enzyme termed uridine phosphorylase (EC: 2.4.2.3) catalyzes the reversible conversion of uridine into uracil [24, 25]. Only recently, two proteins with this enzymatic activity, PcUP1 and PcUP2, have been characterized in the plant pathogenic Oomycete *Phytophthora capsici* [26]. In the next step we looked for putative uridine phosphorylase orthologs using the corresponding *Escherichia coli* protein sequence Udp [27], mammalian UPP1 and UPP2 [28, 29] and that of PcUP1 and PcUP2. However, BLASTP-based analyses did not reveal any putative candidates in *A. fumigatus.* Searching the InterPro database [30, 31] for specific domains within these five proteins, we found that all of them contained a nucleosidase phosphorylase domain, which was also predicted in 10 proteins encoded by the genome of the *A. fumigatus* isolate A1163 (**S2 Table**). This strain is part of the CEA10-lineage, the progenitor of A1160P+ which was used as wt reference in this work [32]. Two out of these proteins, which we termed UdpA (AFUB_097990) and UdpB (AFUB_047570), contained a predicted purine and uridine phosphorylase domain (PNP_UDP_1 domain-containing). To study their potential role in the conversion of 5FU into 5FUR, the respective genes were, like *urhA* and *urhB,* deleted in Δ*uprt.* While no difference in 5FU resistance could be detected between Δ*uprt* and Δ*uprt*Δ*udpA*, a slight increase in resistance was observed for Δ*uprt*Δ*udpB* (**Fig 2A and 2B**). Transforming this double mutant with a functional *udpB* gene cassette (Δ*uprt*Δ*udpB udpB^REC^*), recovered 5FU susceptibility, which further suggests a role of UdpB in the conversion of 5FU into 5FUR. Nevertheless, the small increase in 5FU resistance of Δ*uprt*Δ*udpB* as well as the presence of several further candidate proteins with putative nucleosidase phosphorylase domain indicates that uridine phosphorylase activity is not restricted to a single protein in *A. fumigatus.*

**Fig 2.**
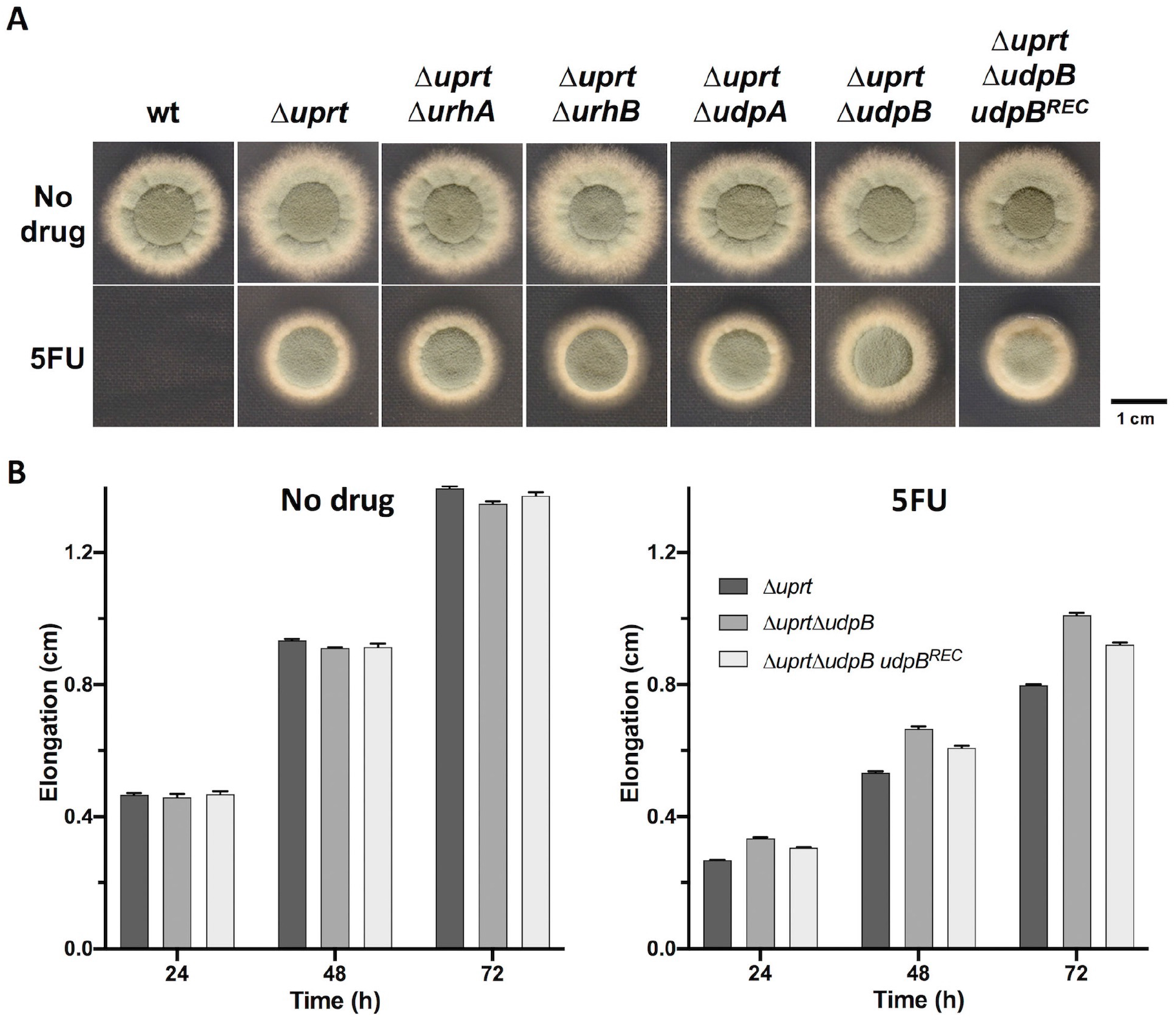
The absence of UdpB increases 5FU resistance in a Uprt deficient background. **(A)** Radial growth of strains lacking Uprt in combination with enzymes with potential uridine phosphorylase activity was analyzed in the presence of 5FU. **(B)** Colony extension rate analyses confirmed the relevance of UdpB in 5FU toxicity. For both experiments, medium was supplemented with 500 μg/mL of 5FU.

### Intracellular metabolization of 5FC precedes efflux of its derivatives into the environment

With the exception of UdpB, we found explicit functions of FcyA, Uprt, UrkA and UrhB within the pyrimidine salvage pathway of *A. fumigatus* that participate in the metabolization of the fluoropyrimidines 5FC, 5FU and/or 5FUR. Specific, high levels of resistance to these compounds were related to the disruption of individual (Δ*fcyA*, Δ*uprt*, Δ*urkA*) or combinations (Δ*urkA*Δ*urhB*, Δ*uprt*Δ*urkA*) of their encoding genes (**Fig 1B**).

The absent salvage activities generating the resistance profiles of these mutants are expected to cause an accumulation of 5FC, 5FU and/or 5FUR within the cell. To study this idea and investigate whether *A. fumigatus* reduces cellular fluoropyrimidine levels by their export into the environment, we measured their intra- and extracellular concentrations in overnight grown liquid cultures that were exposed to 10000 nmol 5FC (100 nmol/mL) for 4 h (**Table 1A** and **1B**).

**Table 1.** Determination of fluoropyrimidine levels in 5FC treated strains. HPLC-based analysis of **(A)** intra- and **(B)** extracellular 5FC, 5FU and 5FUR levels of 5FC treated wt and pyrimidine salvage pathway mutants with highest degrees of resistance to 5FC, 5FU and/or 5FUR; n.d. not detected.

| A | 5FC | 5FU | 5FUR |
| --- | --- | --- | --- |
|  | nmol/mg ± SD |  |  |
| wt | n.d. | n.d. | 0.29 ± 0.02 |
| *ΔfcyA* | 1.14 ± 0.06 | n.d. | n.d. |
| *Δuprt* | n.d. | 0.52 ± 0.06 | n.d. |
| *ΔurkA* | n.d. | 0.32 ± 0.03 | 0.18 ± 0.03 |
| *ΔuprtΔurkA* | n.d. | 0.49 ± 0.06 | n.d. |
| *ΔurkAΔurhB* | n.d. | n.d. | 0.39 ± 0.04 |

| B | 5FC* | 5FU | 5FUR |
| --- | --- | --- | --- |
|  | nmol/mL ± SD |  |  |
|  | 64.3 ± 4.2 | 18.0 ± 1.7 | n.d. |
|  | 85.5 ± 14.8 | n.d. | n.d. |
|  | 35.8 ± 5.0 | 54.8 ± 9.1 | n.d. |
|  | 57.5 ± 1.5 | 25.4 ± 2.4 | 3.4 ± 0.3 |
|  | 26.4 ± 3.2 | 75.6 ± 1.5 | n.d. |
|  | 56.2 ± 2.0 | 19.6 ± 1.6 | 11.9 ± 0.9 |

\*Extracellular 5FC levels constitute a mixture of initially added as well as exported 5FC.

Δ*fcyA* lacking cytosine deaminase activity showed the highest levels of intra- as well as extracellular 5FC. Neither 5FU nor 5FUR could be detected in this mutant. The highest quantities of intra- as well as extracellular 5FU were observed in mutants lacking Uprt (Δ*uprt*, Δ*uprt*Δ*urkA*), which is in agreement with the role of this enzyme catalyzing the main route of 5FU conversion into 5FUMP. Remarkably, in the culture supernatants of Δ*uprt* and Δ*uprt*Δ*urkA*, 54.8 and 75.6 nmol/mL 5FU were detected, respectively. The largest amount of intra- as well as extracellular levels of 5FUR were detected in Δ*urkA*Δ*urhB* lacking uridine kinase and uridine nucleosidase activities. Despite the block in the direct phosphorylation of 5FUR, Δ*urkA* did not show increased intracellular 5FUR levels compared to wt. However, in contrast to wt an increase in extracellular 5FUR could be detected in Δ*urkA,* which is most likely due to an increased export of 5FUR in mutants lacking this enzyme.

### Fluoropyrimidine-efflux during 5FC treatment creates a toxic environment for cytosine deaminase deficient cells

As demonstrated above, during 5FC treatment *A. fumigatus* cells reduce intracellular fluoropyrimidine levels through their export into the environment. Like Δ*fcyA,* mammalian cells lack cytosine deaminase and are therefore resistant to 5FC [11]. Both still have the enzymatic capacity to activate 5FU and 5FUR [17, 20], which suggests that the 5FC derived fluoropyrimidine content in supernatants of 5FC treated strains, particularly that of *uprt* deficient strains, could induce toxic effects on both Δ*fcyA* and mammalian cells. To test this idea, different experimental approaches were conducted.

First, wt, Δ*fcyA, Δuprt* and Δ*uprt*Δ*urkA* were grown on solid *Aspergillus* minimal medium (AMM) in the presence of their 5FC treatment derived culture supernatants (**Fig 3A**). In agreement with Δ*fcyA* not being able to metabolize the supplemented 5FC, its supernatant did not affect its own growth and that of Δ*uprt*Δ*urkA*, which is still capable to convert 5FC into 5FU and 5FUR but not 5FUMP. In line with the presence of the alternative route, growth of Δ*uprt* was slightly impaired by the Δ*fcyA* supernatant with containing high 5FC levels. Δ*uprt* supernatant, comprising high 5FU, led to a clear growth reduction of Δ*fcyA*. The fluoropyrimidine content of this supernatant slightly blocked growth of Δ*uprt* itself and, surprisingly, that of Δ*uprt*Δ*urkA*. Inhibition of the double mutant, which is anticipated to lack 5FUMP formation out of 5FC, 5FU or 5FUR, suggests that additional but so far unknown 5FC derived inhibitory fluoropyrimidines are generated and exported in Δ*uprt*. These toxic metabolites, which could still be produced through the alternative pathway, might consist of 5FUMP and its derived 5-fluorinated nucleotides. Similar to that of Δ*uprt,* supernatant of wt expressing the full set of pyrimidine salvage enzymes, also exhibited low activity against Δ*uprt*Δ*urkA*. Furthermore, this supernatant showed high antifungal activity against Δ*fcyA* and Δ*uprt*, which is most likely a result of exported 5FUR, 5FUMP and its derived 5-fluorinated nucleotides. Supernatant of Δ*uprt*Δ*urkA*, with the remaining enzymatic capacity to produce 5FU and 5FUR, caused the highest growth inhibition of Δ*fcyA* and slightly hampered growth of Δ*uprt.* In accordance with its lack of enzymes to generate 5FUMP, Δ*uprt*Δ*urkA* was not inhibited by its own supernatant.

**Fig 3.**
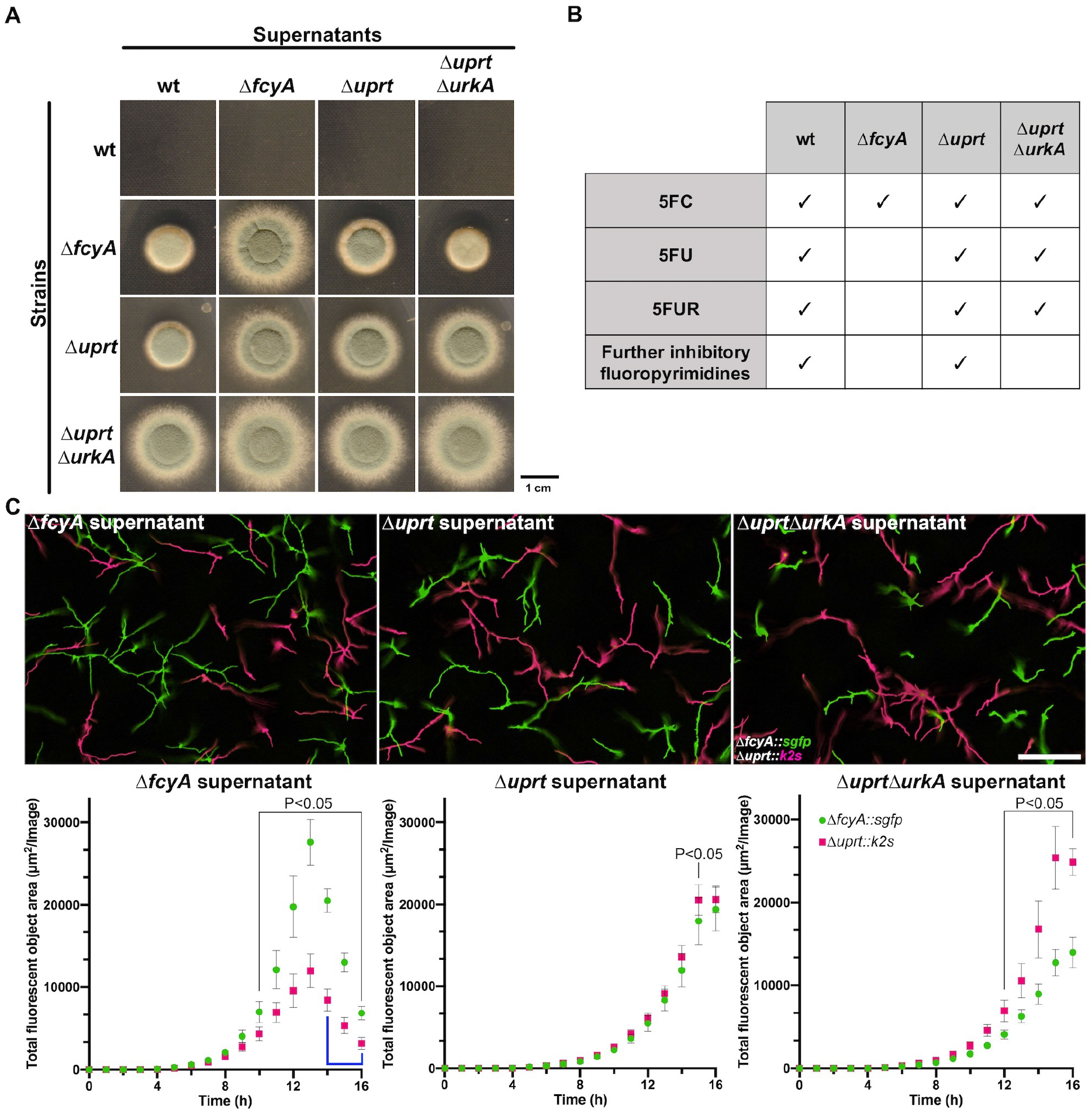
Growth inhibitory effects exerted by effluxed fluoropyrimidines. **(A)** Evaluation of the supernatants impact on the colony growth of mutants lacking key pyrimidine salvage activities. **(B)** Prediction of the fluoropyrimidine content in supernatants of the respective strains. The slight susceptibility displayed by Δ*uprt*Δ*urkA* grown in the presence of wt and Δ*uprt* supernatants suggests the efflux of 5FUMP and/or further toxic 5-fluorinated nucleotides. **(C)** Microtiter-based co-cultivation of Δ*fcyA* and Δ*uprt* expressing sGFP and K2S, respectively. Exemplary micrographs of their hyphal growth after 13 h (Δ*fcyA* supernatant) and 15 h (Δ*urpt* and Δ*urptΔurkA* supernatants) are shown (upper panel). Growth difference between mutants was determined by quantifying the total amount of area within the image containing GFP or RFP fluorescence. After 13h incubation in Δ*fcyA* supernatant, 3-dimensional growth of the mycelium led to autofocus related imprecisions in image acquisition as previously reported [52] (blue line). Scale bar: 200 μm.

In the second approach, we carried out microtiter-based co-cultivation experiments in liquid AMM to simultaneously analyze growth behavior of Δ*fcyA* and Δ*uprt* during supplementation with different culture supernatants (**Fig 3C**). To achieve this, sequences encoding *fcyA* or *uprt* were replaced by expression cassettes which encoded the green fluorescent protein variant GFP^S65T^ [33] and the red fluorescent protein Katushka2S [34], here abbreviated as sGFP and K2S respectively, allowing their individual monitoring within the same well using fluorescence microscopy. Similar to the results obtained on solid AMM, in the presence of Δ*fcyA* supernatant, Δ*fcyA* showed elevated resistance when compared to Δ*uprt.* During incubation in supernatants of Δ*uprt* and Δ*uprt*Δ*urkA*, both with expected high 5FU levels, Δ*uprt* displayed higher growth rates than Δ*fcyA.*

In a third approach, we investigated antiproliferative properties of culture supernatants on the human lung carcinoma epithelial cell line A549 [35]. Therefore, A549 cells were incubated in Dulbecco’s Modified Eagle Medium (DMEM) supplemented supernatants of 5FC treated wt, Δ*fcyA, Δuprt* and Δ*uprtΔurkA.* Supernatant of Δ*fcyA* served as control as it should only contain 5FC and thus not induce fluoropyrimidine related inhibitory effects. In agreement, A549 cells showed comparable growth behavior in medium supplemented with Δ*fcyA* supernatant or water (**Fig 4A** and **4B**). Similar to the results obtained for growth *of A. fumigatus ΔfcyA,* A549 cells showed the largest growth defects when incubated with supernatants of wt and Δ*uprt*Δ*urkA* followed by that of Δ*uprt.* Together, this result leads to the suggestion that fluoropyrimidine efflux by *A. fumigatus* could enhance cytotoxic effects during 5FC antifungal treatment.

**Fig 4.**
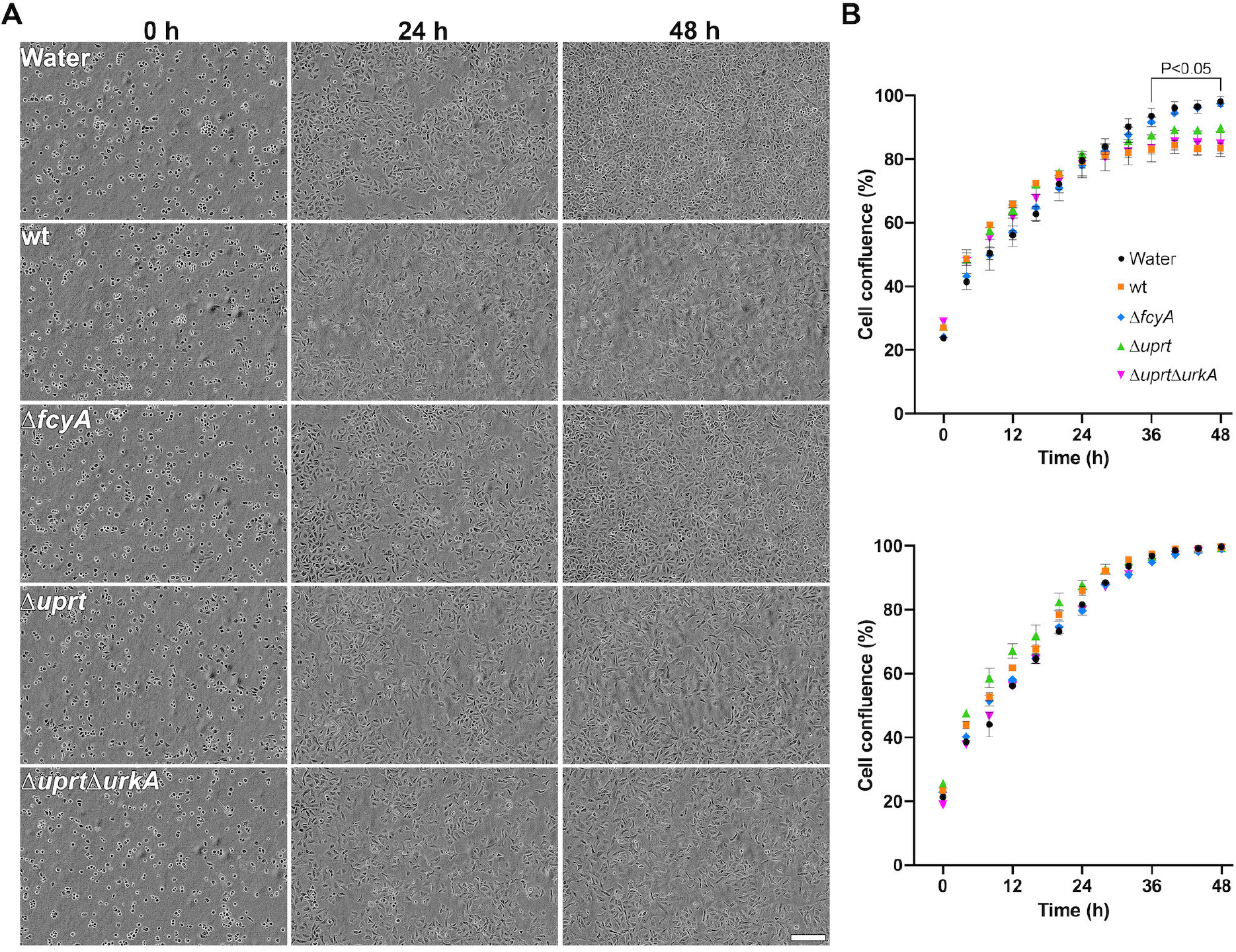
Fungal supernatants containing 5FC-derived antimetabolites compromise proliferation of mammalian cells. **(A)** Microscopy-assisted analysis of A549 cells exposed to 5FC treatment derived fungal culture supernatants. **(B)** Cell confluence was determined over a period of 48 h to monitor fluoropyrimidine based growth inhibition (upper panel). Supernatants of *A. fumigatus* strains that were treated with water instead of 5FC served as control (lower panel). Scale bar: 200 μm.

### Efflux inhibition increases susceptibility to 5-fluoropyrimidines

As stated above, at neutral pH *A. fumigatus* displays intrinsic 5FC resistance due to transcriptional downregulation of *fcyB* encoding the major 5FC import protein in this fungus [14]. Based on our data we suspected efflux-mediated intracellular reduction of fluoropyrimidines during 5FC treatment to be an additional factor contributing to 5FC tolerance that could be overcome by its inhibition. To test this idea, we investigated synergistic interactions of 5FC, 5FU and 5FUR with the fungal ABC and MFS efflux pump inhibitor clorgyline [18, 19], here abbreviated as CLG. For this, checkerboard assays coupled with high-throughput microscopic analysis were employed. Based on the growth inhibitory action of each compound as well as their combinations, Bliss synergy scores were calculated using the SynergyFinder Plus tool [36, 37]. For each 5FC, 5FU and 5FUR, synergistic interaction with CLG was found at various combinations (**Fig 5A** and **5B**, **S2A** and **S2B Fig**, **S3A and S3B Table**). The highest scores were determined using CLG at a concentration of 25 and 50 μg/mL. On their own for instance, visually assessed MIC levels of 5FC and 5FU were 200 and 100 μg/mL in AMM as well as 400 and 100 μg/mL in RPMI, respectively. Despite the obvious antifungal activity of 5FUR, even at a concentration level of 2000 μg/mL wt was still able to grow in both media and therefore, no MIC could be determined for this drug. In combination with 25 μg/mL CLG, we detected MICs for 5FC, 5FU and 5FUR of 12.5, 6.25 and 31.25 μg/mL, respectively, in AMM and 25, 6.25 and 250 μg/mL in RPMI using 50 μg/mL CLG. The large increase of 5FC activity (up to 16-fold) in the presence of CLG indicates fluoropyrimidine export as a mechanism in *A. fumigatus* that significantly lowers 5FC activity, which can be overcome by combinatorial treatment.

**Fig. 5.**
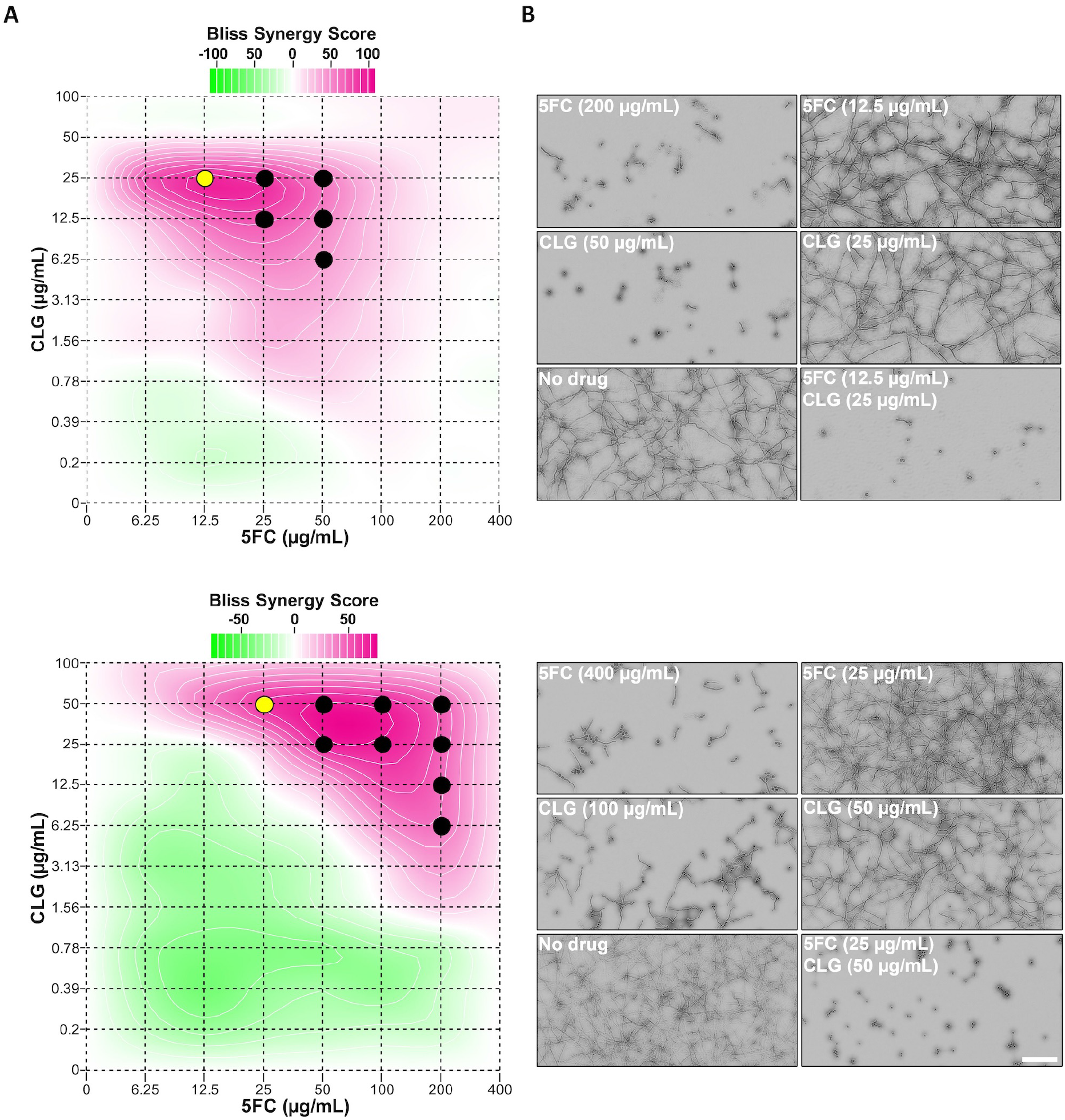
CLG potentiates 5FC antifungal activity against *A. fumigatus.* **(A)** Checkerboard assays using 5FC in combination with CLG were carried out to determine Bliss synergy scores. Circles represent combinations of 5FC and CLG where full growth inhibition was observed. **(B)** Microscopy images of wells corresponding to the visually detected MICs of 5FC and CLG in single use as well as their combination requiring the lowest 5FC amount to prevent fungal growth (yellow circle). Checkerboard assays were performed with AMM (upper panels) and RPMI (lower panels). Scale bar: 100 μm.

## Discussion

5FC metabolic activation starts with the formation of the nucleotide 5FUMP and, in *A. fumigatus*, this process is predominantly mediated by cytosine deaminase (FcyA) and uracil phosphoribosyltransferase (Uprt) [20]. In this work, we identified three further enzymes with pyrimidine salvage activities including uridine phosphorylase (UdpB), uridine nucleosidase (UrhB) and uridine kinase (UrkA), which mediate the metabolization of 5FC derivatives 5FU and/or 5FUR. Two of them, uridine phosphorylase and uridine kinase, enable an alternative enzymatic route of 5FC metabolic activation in which 5FU, the reaction product of cytosine deaminase, is converted to 5FUMP *via* 5FUR (**Fig 6**). Regarding UdpB, our findings suggest that the conversion of 5FU into 5FUR is not solely mediated by this protein but rather several enzymes.

**Fig 6.**
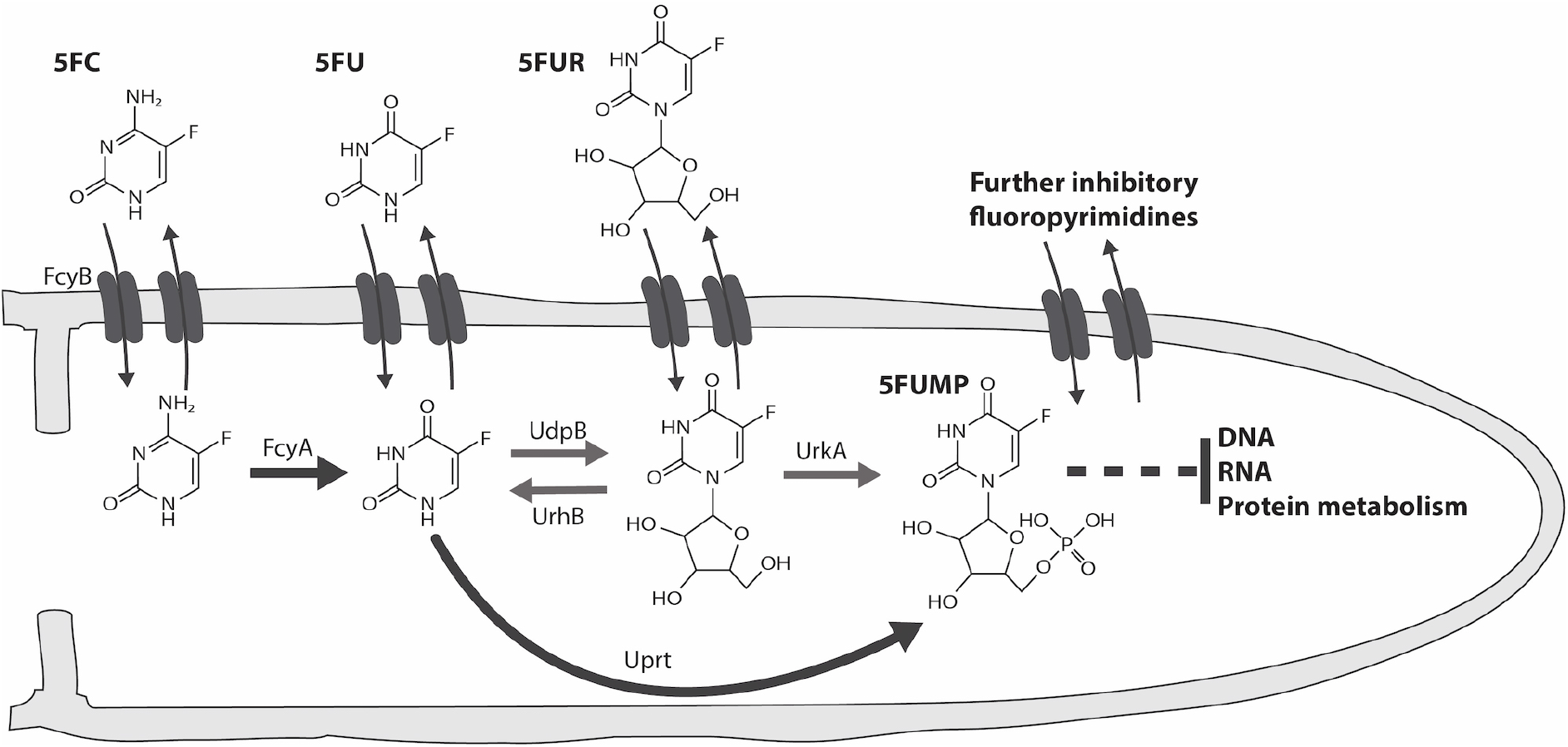
Proposed model illustrating transport dynamics and pyrimidine salvage activities that mediate the metabolic conversion of fluoropyrimidines in *A. fumigatus.* The major route of 5FC metabolic activation is catalyzed by FcyA and Uprt (black arrows). UdpB, UrhB and UrkA constitute enzymes of a metabolic bypass (grey arrows) that generates 5FUMP, however in a less efficient way. During 5FC treatment, intracellular levels of 5FC, 5FU, 5FUR and so far unknown fluoropyrimidine antimetabolites are reduced by their export into the environment.

During 5FC treatment, the metabolic blocks caused by the absence of specific pyrimidine salvage activities resulted in increased intracellular accumulation of either 5FC (Δ*fcyA*), 5FU (Δ*uprt* and Δ*uprt*Δ*urkA*) or 5FUR (Δ*urkA*Δ*urhB*) which was accompanied by their elevated efflux into the environment. Particularly mutants lacking Uprt, catalyzing the major and direct route of 5FUMP formation, showed excessive extracellular 5FU levels in their culture supernatants after 5FC exposure. To further conclude about the effluxed fluoropyrimidine composition of Uprt mutants as well as that of Δ*fcyA* (5FC accumulation control) and wt expressing the full set of pyrimidine salvage enzymes, growth inhibition assays were performed using medium supplemented with their supernatants (**Fig 3**). In this regard it has to be mentioned that in contrast to Δ*uprtΔurkA,* both wt and Δ*uprt* possess the enzymatic capacity to generate 5FUMP, which represents the precursor of various active 5-fluorinated nucleotides. Susceptibility profiles of wt, Δ*fcyA, Δuprt* and Δ*uprt*Δ*urkA* suggested the efflux of 5FU and 5FUR by Δ*uprtΔurkA,* together with so far unidentified toxic fluoropyrimidines by wt and Δ*uprt*.

As stated above, 5FC is considered non-toxic for mammalian cells due to the absence of cytosine deaminase [11]. Yet, cytotoxic effects are occurring during 5FC treatment that were, at least in part, associated with the capacity of members of the human microbiome to convert 5FC into the chemotherapeutic agent 5FU [38, 39]. The composition of microbial species and their expressed salvage activities as well as their location within the body probably constitute several factors that could influence 5FC related host toxicity. *E. coli,* a common member of human microbiome [39], for instance expresses cytosine deaminase [40], uracil phosphoribosyltransferase [41], uridine phosphorylase [27] as well as uridine kinase [42], which should enable the conversion of 5FC into 5FU, 5FUR, 5FUMP as well as further derivatives thereof. Thus, it is very likely that in addition to 5FU, a variety of other 5FC derived cytotoxic compounds will be generated and released by the human microbiome during 5FC exposure. We hypothesized that, in addition to the fluoropyrimidines produced by the microbiome, those produced by pathogens could enhance adverse effects on host cells during infections involving 5FC treatment. Using culture supernatants of 5FC exposed wt and Uprt deficient mutants, in this work we demonstrate the capacity of *A. fumigatus* to generate and release fluoropyrimidines that compromise proliferation of A549 cells. We speculate that similar fluoropyrimidine detoxifying mechanisms than those observed for *A. fumigatus* could be present in clinical yeast, which remains to be investigated. Importantly, mutations affecting uracil phosphoribosyltransferase activity are a major reason for cross-resistance to 5FC and 5FU in *Candida* and *Cryptococcus* species [11].

In previous work we elucidated transcriptional repression of 5FC uptake as the major mechanism conferring high 5FC tolerance in *A. fumigatus* at neutral pH [14]. Here, we uncover fluoropyrimidine efflux as an additional factor that diminishes 5FC activity in this mold pathogen. Based on the hypothesis that 5FC activity could be enhanced by perturbing fluoropyrimidine export, we employed 5FC as well as its effluxed derivatives 5FU and 5FUR in combination with the fungal efflux pump inhibitor CLG and identified strong synergistic interactions between CLG and all three compounds. The fluoropyrimidine export-based adverse effects on 5FC antifungal activity as well as host cell proliferation highlight fungal efflux as an attractive therapeutic target during 5FC treatment.

## Material and methods

### Growth conditions for phenotypic analyses and fungal transformation

Generally, AMM [43] containing 20 mM ammonium tartrate as nitrogen source and 1% glucose as carbon source was used for phenotypic analysis. Plate growth assays were conducted on solid AMM containing 1.5% agar (Carl Roth GmbH, Karlsruhe, Germany). Cultures were generally grown at 37 °C.

For plate growth-based susceptibility testing of mutants, 10^4^ spores in a total volume of 5 μL spore buffer (0.1% Tween 20, 0.9% sodium chloride solution *(w/v))* were point inoculated on solid AMM supplemented with 5FC, 5FU or 5FUR (Tokyo Chemical Industry, Tokyo, Japan). Colony extension measurements were performed by inoculating the respective strain on the edge of AMM plates. Subsequently, the mean of the colony extension rate was calculated after measuring five vertical transects on the edge of the colony coverage area every 24 h for a final period of three days.

For fungal transformation, solid AMM with 1 M sucrose and 0.7% agar was used. Solid Sabouraud dextrose (SAB, Sigma-Aldrich Corp., St. Louis, MI, USA) medium with 1.5% agar was used for the generation of spores. Fungal genetic manipulations were carried out using 1 μg of the corresponding DNA construct which was transformed into the respective recipients. Selection procedures using conventional selectable marker genes as well as counterselectable markers *fcyA* and *uprt* were conducted as described previously for *A. fumigatus* [20, 44].

### Deletion of *A. fumigatus urkA, urhA, urhB, udpA* and *udpB*

The strains and primers used in this study are listed in **S4** and **S5 Table**, respectively. Generally, the CEA10 Δ*ku80* derivative A1160P+, here termed wt, was used as parental strain. To generate knock-out mutants, the coding sequence of the corresponding gene was disrupted using hygromycin B, phleomycin and pyrithiamine resistance cassettes. Therefore, deletion constructs comprising approximately 1 kb of the respective 5’ and 3’ nontranslated regions (NTRs) linked to the central antibiotic resistance cassette were generated using fusion PCR as previously described [45]. Correct integration of transformed constructs was confirmed by Southern blot analysis.

### Reconstitution of the Δ*urkA, ΔurhB* and Δ*udpB* mutants

To complement the Δ*urkA, ΔurhB and ΔudpB* mutant phenotypes, plasmids pFG66, pFG67 and pESV38 were generated (**S6 Table**). The strategy used to complement mutants lacking different pyrimidine salvage activities is illustrated in **S3 Fig**. First, the coding sequences plus approximately 1-kb 5’ and 3’ of their respective NTRs were amplified using primers urkAcompl-FW/-RV, urhBcompl-FW/RV and udpBcompl-FW/RV. Second, the backbone of the pyrithiamine resistance-conferring plasmid pSK275 [46] was amplified with the primer pair BBpSK275-FW/RV. After PCR-purification, all PCR products were assembled into the pSK275 backbone using the NEBuilder HiFi DNA assembly Master Mix (New England Biolabs, Ipswich, MA, USA), and the constructs were propagated in *E. coli*. Around 1 *μ*g of the pFG66, pFG67 and pESV38 plasmids were linearized by restriction digestion with *Eco*RV, *HpaI* and *MluI* respectively and used as templates to transform into the corresponding genetic background.

### Generation of Δ*fcyA* and Δ*uprt* strains expressing fluorescent proteins

The strategy for the generation of plasmids carrying sequences encoding sGFP and K2S is represented in **S4 Fig**. A common backbone was amplified using the plasmid pAN7-1 [47] as DNA template for the primer set BBgpdA-FW/RV. The inserts encoding sGFP and K2S were amplified using pgfpcccA [48] and a synthetic gene strand (Integrated DNA Technologies, Coralville, IA, USA; codon adapted for *A. fumigatus),* with the primer pairs PgpdAsGFP-FW/RV and PgpdAK2S-FW/RV, respectively. The assembled plasmids, termed pFG36 and pFG39, were used as DNA template for the primer set hph-FW/RV to generate the sGFP and K2S expression cassettes under the control of the *A. nidulans gpdA* promoter (*PgpdA*). Fragments for homologous recombination driven exchange of these two markers were generated and transformed similarly to the procedure described previously [20]. Approximately 1-kb 5’ and 3’ NTRs of *fcyA* and *uprt* were linked to the respective expression cassette using fusion PCR. Subsequently, the knock-in constructs were transformed into wt, generating the Δ*fcyA::sgfp* and Δ*uprt::k2s* fluorescent strains.

### Efflux-based growth inhibition and promotion assays on solid AMM

Supernatants from wt, Δ*fcyA*, Δ*uprt* and Δ*uprt*Δ*urkA* grown in the presence of 5FC were recovered. For this, 100 mL of liquid AMM was inoculated with 1×10^6^/mL spores of each strain and incubated at 37 °C for 16 h shaking at 200 rpm. Cultures were then supplemented with 250 μg/mL of 5FC and incubated for further 10 h before collecting. Before sterile filtration, the supernatants were incubated at 80 °C for 1 h to prevent metabolic activities. At last, 10 mL 2× AMM containing 3% agar was combined with 10 mL of each supernatant for plate growth-based susceptibility testing.

### Microscopic analysis of efflux-based growth inhibition

For microscopic monitoring of inhibitory effects of effluxed 5-fluoropyrimidines, Δ*fcyA::sgfp* and Δ*uprt::k2s* fluorescent mutants were co-inoculated in liquid AMM in Nunc96 microplates (Thermo Scientific Inc., Waltham, MA, USA). For each strain 1×10^4^ spores in a total volume of 25 μL 2× AMM were inoculated in each well. Then, 50 μL of the supernatants generated by different salvage mutants were added giving a final volume of 100 μL. The microplates were incubated at 37 °C for 16 h and monitored with the IncuCyte S3 Live-Cell Analysis System equipped with a 20× magnification S3/SX1 G/R Optical Module (Essen BioScience Inc., Ann Arbor, MI, USA). Four images per well were acquired every hour. Fungal growth was determined using the Basic Analyzer tool of the IncuCyte S3 software (version 2020B) for fluorescent imaging (GCU 1.400 and RCU 2.000). The background of the fluorescent images was subtracted using ImageJ/FIJI (version 2.3.0/1.53f) to facilitate the visualization of the cells.

### Cell culture and proliferation assay

The human lung carcinoma epithelial cell line A549 (American type culture collection, CCL-185) was maintained by serial passage in DMEM/F-12 (Sigma-Aldrich, St. Louis, MI, USA) with 10% FBS (Sigma-Aldrich, St. Louis, MI, USA). For the proliferation assay, cells were seeded at a concentration of 1×10^5^ cells/mL in Nunc96 microplates and cultured at 37 °C and 5% CO_2_ until a confluent monolayer was formed. Cells were then challenged with 5 μL supernatant (1:20 dilution) collected from 5FC exposed wt, Δ*fcyA, Δuprt* and Δ*uprt*Δ*urkA* liquid cultures. Growth of A549 cells was monitored over the duration of 48 h with the IncuCyte S3 Live-Cell Analysis System. Four images per well were acquired every two h. For A549 cells, the magnification required with the S3/SX1 G/R Optical Module was 10×. Cell confluence was determined using the Basic Analyzer software of the IncuCyte S3 for phase imaging.

### Checkerboard assays and synergy analyses

To study the relevance of efflux pumps on fluoropyrimidine resistance, drug interaction assays were performed by the microdilution checkerboard method [49]. Checkerboards were performed in AMM and RPMI medium (Sigma-Aldrich, St. Louis, MI, USA), combining 5FC, 5FU or 5FUR with clorgyline (CLG) (Sigma-Aldrich, St. Louis MI, USA) against *A. fumigatus* wt. The final concentration of 5FC and 5FU ranged from 6.25 to 400 μg/mL, whereas 5FUR and CLG concentrations ranged from 31.25 to 2000 μg/mL and 0.20 to 100 μg/mL, respectively. In brief, 10 μL of each compound dilution was mixed with 80 μL of 1.25 × AMM or RPMI medium containing 1.25 ×10^5^ spores/mL, to obtain a final volume of 100 μL of 1× medium with 1 *×*10^5^ spores/mL. Microscopic images were acquired after 24 and 48 h of incubation at 37 °C with the IncuCyte S3 Live-Cell Analysis System using the 20× magnification S3/SX1 G/R Optical Module. From each well, two images were taken from the central area of the well. Fungal growth was analyzed using the Basic Analyzer software of the IncuCyte S3 (Confluence %, Segmentation adjustment: 0.1, Adjust size: −1). The Bliss Independence Model [36] was used to analyze the potential synergistic interactions among the different drug combinations. In this model antagonistic interactions are estimated to produce scores less than 10, whereas scores from −10 to 10 are consequence of additive interactions. Scores larger than 10 suggest synergistic interactions.

### Fluoropyrimidine quantification

Intra- and extracellular fluoropyrimidine levels of pyrimidine salvage mutants were quantified by High Performance Liquid Chromatography coupled with Variable Wavelength Detector (HPLC-VWD). For this, 1 × 10^6^ spores/mL of each strain were inoculated in 100 mL of AMM for 16 h. Cultures were then supplemented with 100 nmol/mL of 5FC for further 4 h. Mycelium and supernatant 5FC treated cultures were collected by filtration. Both supernatant and mycelia samples were incubated in a water bath at 80 °C for 1 h before analysis.

Then, 10 mg of dried mycelium were powderized, mixed with 400 μL water and vortexed with two glass beads for 15 min. The mixture was centrifuged for 10 min at 10,000 g and the supernatant was analyzed by HPLC-VWD using an Agilent Series 1100 HPLC system (Agilent Technologies Deutschland GmbH, Waldbronn, Germany). Chromatographic separation was carried out with an Agilent Zorbax Eclipse Plus C18 (150 × 4.6 mm, i.d. 5.0 μm) column (Agilent Technologies Deutschland GmbH, Waldbronn, Germany). The mobile phase was a mixture of water with 0.0001% 1-octanesulfonic acid sodium salt *(w/v)* [50] (A) and methanol (B). The gradient started with 1% (B), followed by 2.5% at 4 min, and by 15% at 4 min. The total run time including column equilibration was 15 min with a flow rate at 1.2 mL/min, and an injection volume of 10 μL. The column oven was set at 22 °C. The UV detection wavelength was set at 280 nm. Data analysis and instrument control was carried out with Thermo Scientific™ Dionex™ Chromeleon™ 7.2 Chromatography Data System (Thermo Fisher Scientific, Dreieich, Germany). The retention times were 2.4 min for 5FC, 2.9 min for 5FU, and 7.2 min for 5FUR. The concentration of each compound was determined using external standard calibration.

### Statistical analysis

GraphPad Prism 9 software (Dotmatics, Boston, MA, USA) was used to analyze and display statistic results. All experiments were performed using three biological replicates. Experiments considering two independent variables (*Aspergillus fumigatus* strains and/or A549 cell line) were analyzed using a two-way ANOVA with Šídák or Dunnett multiple comparison tests. All graphs depict the mean, and the error bars show the standard deviation. *P*-values of ≤0.05 were considered as significant.

## Acknowledgments

This research was funded by the Austrian Science Fund (FWF), grant P31093 to F.G. and M2867 to C.B. L.E. S.-V. participated in the HOROS program W1253 funded by the FWF. The authors would like to thank Johanna Gostner for supplying A549 cells and F2G for providing olorofim. Anna Niedrig and Christoph Müller thank Prof. Dr. Franz Bracher for providing his laboratories and equipment.

