## Supporting information for "The fungal expel of 5-fluorocytosine derived fluoropyrimidines mitigates its antifungal activity and generates a cytotoxic environment"

**S1 Table. Protein-based BLAST analyses (https://blast.ncbi.nlm.nih.gov/Blast.cgi) of *S. cerevisiae* (S288C) uridine kinase Urk1p (YNR012W) and uridine nucleosidase Urh1p (YDR400W) against *A. fumigatus* (A1163). (A)** BLASTP analysis suggested one Urk1p ortholog in *A. fumigatus*. **(B)** Two orthologs of *S. cerevisiae* Urh1p were predicted, which we termed UrhA and UrhB. Despite the higher homology, resistance analysis (**Fig 1** and **S1 Fig**) confirmed uridine nucleosidase activity for UrhB but not UrhA.

**A**

| Description | Score | E-value | Length (aa) | QC | % Identity | Gene ID | Termed |
|---|---|---|---|---|---|---|---|
| Uridine kinase, putative | 338 | 4e-112 | 453 | 90% | 41.81% | AFUB_022460 | UrkA |

**B**

| Description | Score | E-value | Length (aa) | QC | % Identity | Gene ID | Termed |
|---|---|---|---|---|---|---|---|
| Uridine nucleosidase Urh1, putative | 308 | 5e-104 | 358 | 93% | 33.52% | AFUB_005870 | UrhA |
| Nucleoside hydrolase, putative | 169 | 2e-48 | 522 | 88% | 17% | AFUB_011230 | UrhB |

**S2 Table. Domain analyses performed with the InterPro database (https://www.ebi.ac.uk/interpro/) revealed 10 proteins carrying a predicted nucleoside phosphorylase domain (IPR000845) in *A. fumigatus* A1163.** Proteins with a putative purine nucleoside phosphorylase and uridine phosphorylase domains (PNP_UDP_1; PF01048) are highlighted in bold.

| Gene ID | Accession Uniprot | Description Interpro |
|---|---|---|
| AFUB_017990 | B0XT47 | TPR_REGION domain-containing protein |
| AFUB_033090 | B0XVN7 | Pfs and NB-ARC domain protein |
| **AFUB_047570** | **B0XX34** | **PNP_UDP_1 domain-containing protein** |
| AFUB_074680 | B0Y7R0 | S-methyl-5'-thioadenosine phosphorylase |
| AFUB_079240 | B0Y910 | ANK_REP_REGION domain-containing protein |
| AFUB_091840 | B0YB77 | ANK_REP_REGION domain-containing protein |
| AFUB_092960 | B0YCR8 | Purine nucleoside phosphorylase |
| AFUB_097420 | B0YE07 | Pfs, NACHT and Ankyrin domain protein |
| AFUB_097850 | B0YE50 | ANK_REP_REGION domain-containing protein |
| **AFUB_097990** | **B0YE64** | **PNP_UDP_1 domain-containing protein** |

**S3 Tables. CLG exerts synergistic interaction with each 5FC, 5FU or 5FUR.** The Bliss score was determined to assess the potential synergism between CLG and fluoropyrimidines. Checkerboard assays were performed in AMM **(A)** and RPMI **(B)**.

**A**

| AMM | | | | | | | | |
|---|---|---|---|---|---|---|---|---|
| **5FC** | **CLG** | **Score** | **5FU** | **CLG** | **Score** | **5FUR** | **CLG** | **Score** |
| 12.5 | 25 | **96.62** | 25 | 25 | **112.38** | 62.5 | 25 | **43.27** |
| 25 | 25 | **87.21** | 25 | 12.5 | **108.13** | 1000 | 25 | **42.9** |
| 6.25 | 25 | **73.9** | 25 | 6.25 | **105.11** | 125 | 25 | **41.38** |
| 25 | 12.5 | **72.74** | 25 | 3.13 | **103.31** | 31.25 | 25 | **41.22** |
| 12.5 | 12.5 | **63.01** | 12.5 | 25 | **99.04** | 250 | 25 | **40.33** |
| 50 | 25 | **57.23** | 12.5 | 12.5 | **93.54** | 1000 | 12.5 | **38.45** |
| 25 | 6.25 | **53.18** | 6.25 | 25 | **93.06** | 500 | 25 | **37.01** |
| 50 | 12.5 | **52.74** | 25 | 1.56 | **90.75** | 500 | 12.5 | **32.15** |
| 50 | 6.25 | **42.02** | 12.5 | 6.25 | **84.51** | 125 | 12.5 | **31.92** |
| 12.5 | 6.25 | **38.17** | 6.25 | 12.5 | **70.66** | 250 | 12.5 | **31.18** |

**B**

| RPMI | | | | | | | | |
|---|---|---|---|---|---|---|---|---|
| **5FC** | **CLG** | **Score** | **5FU** | **CLG** | **Score** | **5FUR** | **CLG** | **Score** |
| 50 | 50 | **81.77** | 12.5 | 25 | **87.02** | 500 | 50 | **62.73** |
| 100 | 25 | **77.49** | 12.5 | 50 | **84.85** | 250 | 50 | **58.37** |
| 100 | 50 | **77.23** | 25 | 25 | **68.98** | 31.25 | 50 | **52.85** |
| 50 | 25 | **76.18** | 25 | 50 | **65.59** | 2000 | 50 | **50.46** |
| 25 | 50 | **73.53** | 25 | 12.5 | **57.84** | 125 | 50 | **43.26** |
| 100 | 12.5 | **61.34** | 6.25 | 50 | **57.03** | 62.5 | 50 | **43.18** |
| 200 | 25 | **56.41** | 12.5 | 12.5 | **48.88** | 1000 | 50 | **41.82** |
| 200 | 50 | **55.77** | 50 | 25 | **46.10** | 2000 | 25 | **41.17** |
| 200 | 12.5 | **52.58** | 50 | 50 | **43.85** | 31.25 | 100 | **19.45** |
| 200 | 6.25 | **48.94** | 6.25 | 25 | **42.54** | 500 | 100 | **18.21** |

**S4 Table. Strains used in this study.**

| Strain | Genotype | Reference |
|---|---|---|
| A1160P+ (wt) | Δ*ku80::pyrG* | (1) |
| Δ*fcyA* | Δ*fcyA::hph* | (2) |
| Δ*uprt* | Δ*uprt::ble* | (2) |
| Δ*urkA* | Δ*urkA::hph* | This study |
| Δ*urhA* | Δ*urhA::hph* | This study |
| Δ*urhB* | Δ*urhB::hph* | This study |
| Δ*urkA*Δ*urhA* | Δ*urkA::hph*Δ*urhA::ble* | This study |
| Δ*urkA*Δ*urhB* | Δ*urkA::hph*Δ*urhB::ble* | This study |
| Δ*urkA urkA*$^{REC}$ | Δ*urkA::hph urkA*$^{REC}$, *ptrA* | This study |
| Δ*urkA*Δ*urhB urhB*$^{REC}$ | Δ*urkA::hph*Δ*urhB::ble urhB*$^{REC}$, *ptrA* | This study |
| Δ*uprt*Δ*urkA* | Δ*uprt::ble*Δ*urkA::hph* | This study |
| Δ*uprt*Δ*urhA* | Δ*uprt::ble*Δ*urhA::hph* | This study |
| Δ*uprt*Δ*urhB* | Δ*uprt::ble*Δ*urhB::hph* | This study |
| Δ*uprt*Δ*udpA* | Δ*uprt::ble*Δ*udpA::ptrA* | This study |
| Δ*uprt*Δ*udpB* | Δ*uprt::ble*Δ*udpB::hph* | This study |
| Δ*uprt*Δ*udpB udpB*$^{REC}$ | Δ*uprt::ble*Δ*udpB::hph udpB*$^{REC}$, *ptrA* | This study |
| Δ*fcyA::sgfp* | Δ*fcyA::PgpdA-sgfp* | This study |
| Δ*uprt::k2s* | Δ*uprt::PgpdA-katushka2s* | This study |

**S5 Table. Oligonucleotides used in this study.**

| Primer set | Forward primer (5´ → 3´) | Reverse primer (5´ → 3´) | PCR product |
|---|---|---|---|
| fcyA-1/2 | TTGAAACTCCGAGGAAGTCG | TAGTTCTGTTACCGAGCCGGTATGTGGATCCAGAGCGTCA | 5´ fcyA |
| fcyA-3/4 | GCTCTGAACGATATGCTCCCTTCGACAAAATGCCATTGAA | TACCTCCCCGAATACCATGA | 3´ fcyA |
| fcyA-N1/N2 | CGAGTCGCCTTAAAATGAGC | GTGGATCGGTATGCAGGATT | *fcyA* knock-in construct |
| uprt-1/2 | GGAAGGACAGGTACGCCATA | TAGTTCTGTTACCGAGCCGGCGGAGCACTCTGAAAATTGG | 5´ uprt |
| uprt-3/4 | GCTCTGAACGATATGCTCCCTCCCATCGTGTAGCGACATA | TACTACCTTCGCCCTCTGGA | 3´ uprt |
| uprt-N1/N2 | TTTGAGCGATTAAGGTGCAA | GCCCCACTACTTGTTTCCAG | u*prt* knock-in construct |
| urkA-1/2 | ATAGGTGGTAGGGCAGGAGG | TAGTTCTGTTACCGAGCCGGATTAGAATGCGGCGCAACAG | 5´ urkA |
| urkA-3/4 | GCTCTGAACGATATGCTCCCGGTCTATAGTGTCAGGCGGC | TACTACCTTCGCCCTCTGGA | 3´ urkA |
| urkA-N1/N2 | GCCAGAATGAATCGCAGTGC | TGCGATTCGTGACTTCTCCC | *urkA* deletion constructs |
| urhA-1/2 | TCACGCAATCTCTGCTCAAG | TAGTTCTGTTACCGAGCCGGGCCAGAATCCTCATGAAACC | 5´ urhA |
| urhA-3/4 | GCTCTGAACGATATGCTCCCGAACCTGGCGATCATAGACC | ACTGATCGGCTGACGTTTTT | 3´ urhA |
| urhA-N1/N2 | GGAGGCAACAATTCTTCCAA | TCACCAAGTGTGCTCGACTC | *urhA* deletion constructs |
| urhB-1/2 | AACTGAAGGAATCGCTGGAA | TAGTTCTGTTACCGAGCCGGTCGCATTACCCGTTTACTCC | 5´ urhB |
| urhB-3/4 | GCTCTGAACGATATGCTCCCAAGCATGCGCCTTTCATTAG | GAACGGGTCAATTGCGTATT | 3´ urhB |
| urhB-N1/N2 | CGGAGTAGCACTGGGAAGTC | GTGCTGATAGCGGAAGGAAG | *urhB* deletion constructs |
| udpA-1/2 | GGATATCGATCCGACTCTCG | TAGTTCTGTTACCGAGCCGGACGAGGCCAACAACAACAA | 5´ udpA |
| udpA-3/4 | GCTCTGAACGATATGCTCCCCTTCTCGTGGAAGTGGTACTG | TCCGAACTGAAAGCCTTTGT | 3´ udpA |
| udpA-N1/N2 | TTCCAAACCCTAATGCCAAG | TGCTGGTCTGACAATCGAAG | *udpA* deletion constructs |
| udpB-1/2 | GTGGTACGGCAGTGAGCAGT | TAGTTCTGTTACCGAGCCGGACAAATCGTGGGAACGAGAC | 5´ udpB |
| udpB-3/4 | GCTCTGAACGATATGCTCCCGCATCCCTTGCATCCATAGT | CAAGTTCAGCAAGGGGGTTA | 3´ udpB |
| udpB-N1/N2 | TTTATGGCCCGATTGCTTAG | GGGGCTCGGTCATAGTAGGT | *udpB* deletion constructs |
| urkAcompl-FW/RV | CCGGCTCGGTAACAGAACTATCCCACCGAGTGTATGATGA | GGGAGCATATCGTTCAGAGCCTTGTTCGTCGCGAAAATCT | Insert for pSK275 backbone (*urkA*) |
| urhBcompl-FW/RV | CCGGCTCGGTAACAGAACTAAGCGACCAGCTCTTTACCAA | GGGAGCATATCGTTCAGAGCCCCGAAGAAAAAGAGACAGC | Insert for pSK275 backbone (*urhB*) |
| udpBcompl-FW/RV | CCGGCTCGGTAACAGAACTACCCAGAAGACGGCCTATGAT | GGGAGCATATCGTTCAGAGCCGGTATAGTCCTCTGGGCAG | Insert for pSK275 backbone (*udpB*) |
| BBpSK275-FW/RV | GCTCTGAACGATATGCTCCCGCTTATCGATACCGTCGACCT | TAGTTCTGTTACCGAGCCGGAATGCCCCACCGTTACATAC | pSK275 backbone for genetic complementations |
| PgpdAsGFP-FW/RV | CCGCTTGAGCAGACATCACCATGGTGAGCAAGGGCGAG | TCCCGCGGTCGGCATCTACTTCACTTGTACAGCTCGTCCA | Insert for pAN7-1 backbone (*sgfp*) |
| PgpdAK2S-FW/RV | CCGCTTGAGCAGACATCACCATGGTCGGCGAGGACTCCGT | TCCCGCGGTCGGCATCTACTCTAGGAGTGGCCCAGCTTGG | Insert for pAN7-1 backbone (*katushka2s*) |
| BBgpdA-FW/RV | AGTAGATGCCGACCGCGG | GGTGATGTCTGCTCAAGCG | pAN7-1 backbone for *sgfp* or *katushka2s* inserts |
| hph-FW/RV | CCGGCTCGGTAACAGAACTAACGGCGTAACCAAAAGTCAC | GGGAGCATATCGTTCAGAGCTCTTGACGACCGTTGATCTG | *hph*, *ble*, *PgpdA-sGFP* and *PgpdA-K2S* cassettes |
| ptrA 5/3 | CCGGCTCGGTAACAGAACTAGCATCCCATTGGTAACGAAA | GGGAGCATATCGTTCAGAGCAATGCCCCACCGTTACATAC | *ptrA* cassette |

**S6 Table. Plasmids used in this study.**

| Plasmid | Used for | Reference |
|---|---|---|
| pAN7-1 | *hph* cassette amplification | (3) |
| pAN8-1 | *ble* cassette amplification | (4) |
| pSK275 | Amplification of pSK275-backbone and *ptrA* cassette | (5) |
| pgfpcccA | *gfp*$^{S65T}$ (here termed as sGFP) amplification | (6) |
| pFG36 | *PgpdA-sgfp* cassette amplification | This study |
| pFG39 | *PgpdA-k2s* cassette amplification | This study |
| pFG66 | *urkA* reconstitution | This study |
| pFG67 | *urhB* reconstitution | This study |
| pESV38 | *udpB* reconstitution | This study |

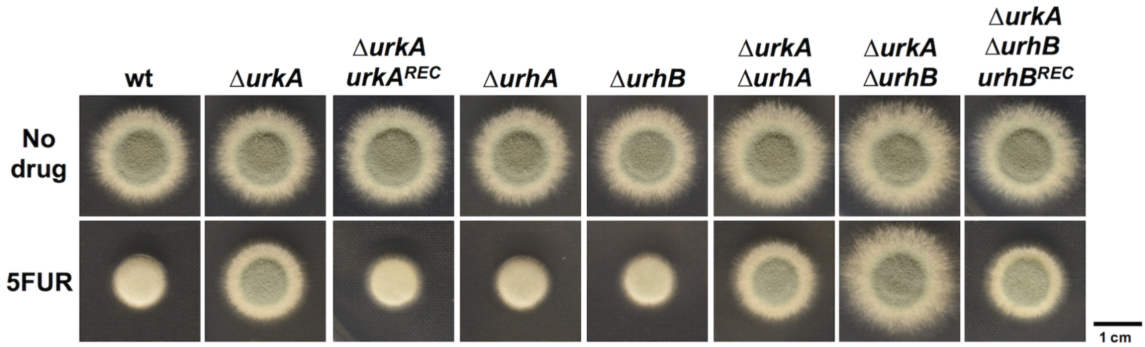


**S1 Fig. Plate growth-based susceptibility testing of *urkA*, *urhA* and *urhB* mutants with predicted defects in 5FUR/uridine metabolization.** 5FUR resistance of deletion mutants as well as complemented versions (*REC*) was assessed on solid medium supplemented with 500 µg/mL of 5FUR.

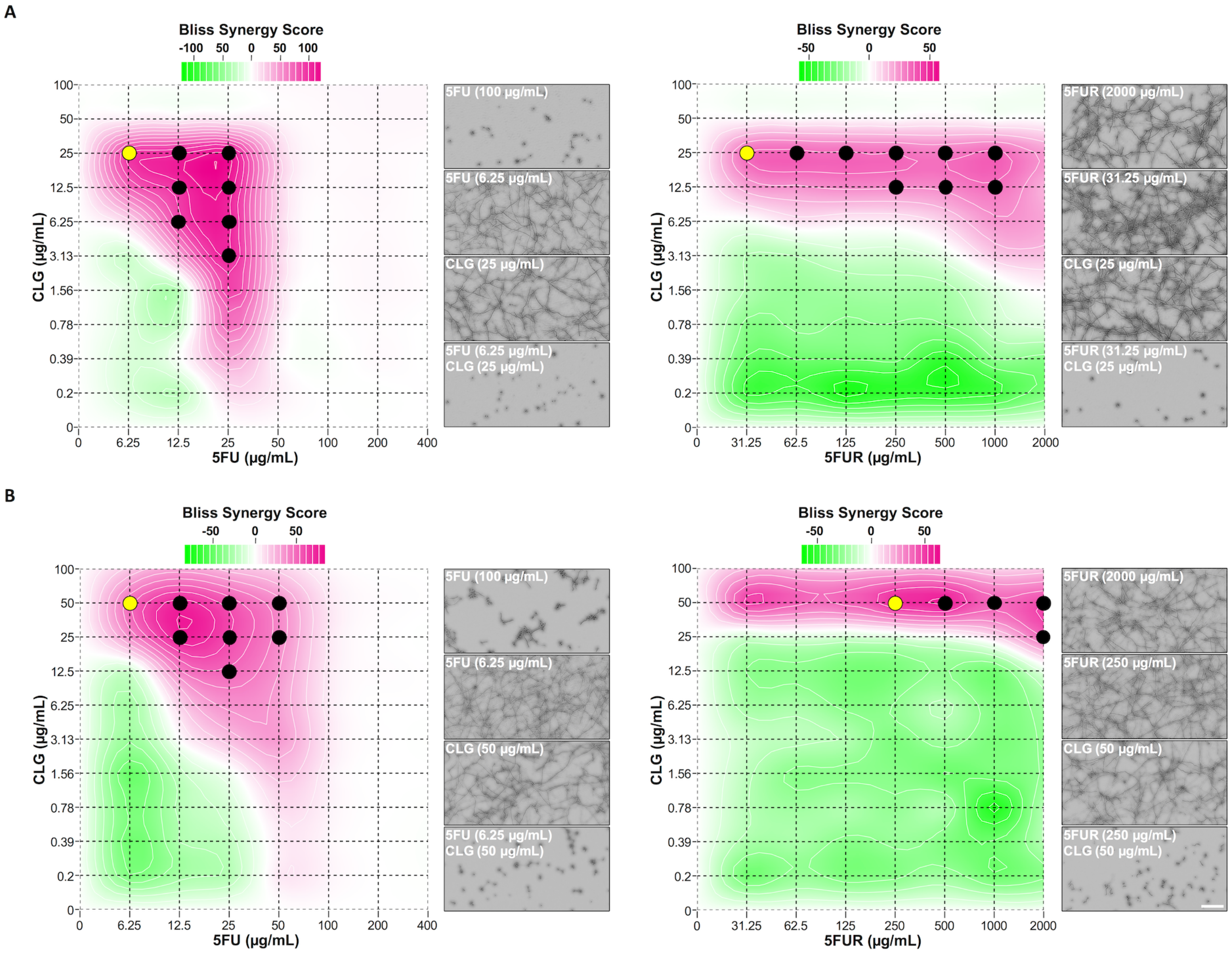


**S2 Fig. Combinations of both 5FU and 5FUR with CLG show strong synergistic effects.** Synergistic interaction between the compounds was analyzed in AMM **(A)** and RPMI **(B)**. Circles represent drug combinations displaying full growth inhibition. Microscopic images of wells corresponding to the combination for which the lowest amount of fluoropyrimidines (yellow circles) were required to prevent growth. In addition, images of wells corresponding to the visually detected MIC of 5FU as well as the maximum amount of 5FUR in single use are illustrated. Scale bar: 100 µm.

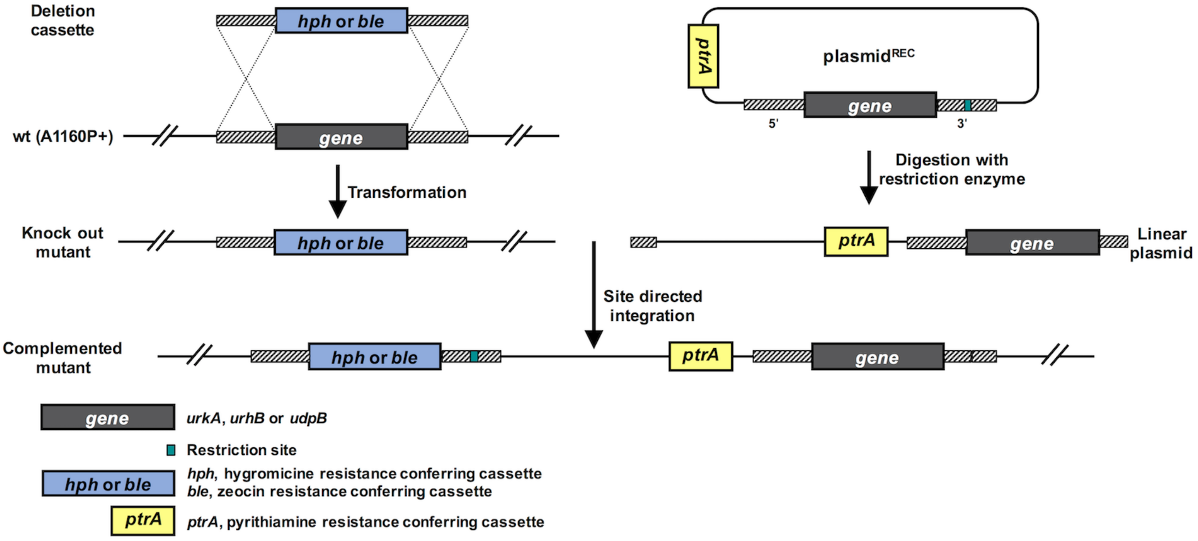


**S3 Fig. Schematic illustration of the strategy used to generate pyrimidine-salvage knock out and complemented mutants.** Plasmid for the reconstitution were linearized for site-directed insertion at the corresponding deletion locus.

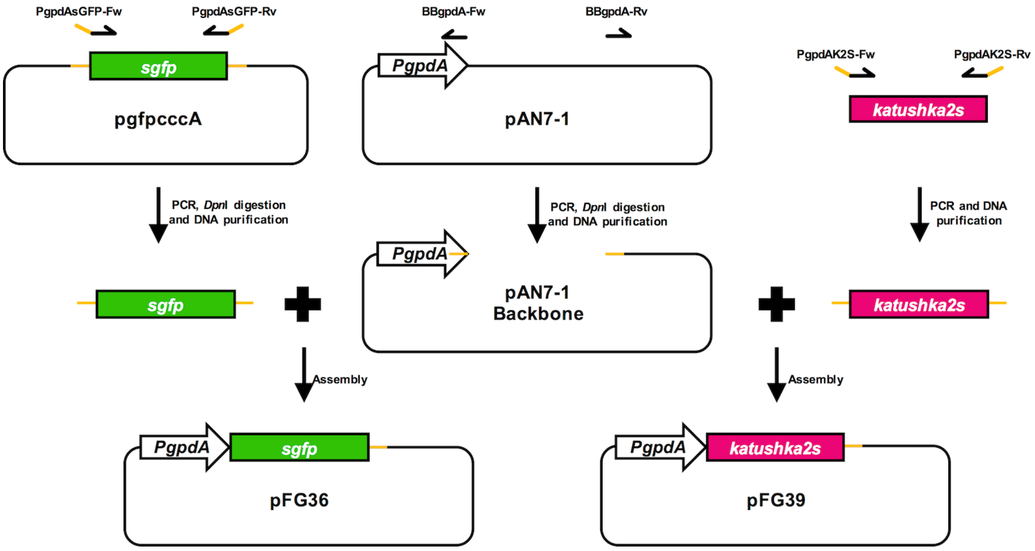


**S4 Fig. Scheme representing the generation of plasmids with fluorescent reporter encoding genes**.
